# Does the impact of cultivar mixtures on virulence dynamics in *Zymoseptoria tritici* populations persist after interseason sexual reproduction?

**DOI:** 10.1101/2022.01.24.477584

**Authors:** Carolina Orellana-Torrejon, Tiphaine Vidal, Sébastien Saint-Jean, Frédéric Suffert

**Affiliations:** Université Paris-Saclay, INRAE, UR BIOGER, 78850 Thiverval-Grignon, France; Université Paris-Saclay, INRAE, AgroParisTech, UMR ECOSYS, 78850 Thiverval-Grignon, France

**Keywords:** Cultivar mixtures, interseason, Septoria leaf blotch, sexual reproduction, *Stb16q*, virulence, wheat disease epidemics, *Zymoseptoria tritici*

## Abstract

This study follows on from a previous study showing that binary mixtures of wheat cultivars affect the evolution of *Zymoseptoria tritici* populations within a field epidemic from the beginning (t1) to the end (t2) of a growing season. Here, we focused on the impact of interseason sexual reproduction on this evolution. We studied mixtures of susceptible and resistant cultivars (carrying *Stb16q*, a recently broken-down resistance gene) in proportions of 0.25, 0.5 and 0.75, and their pure stands. We determined the virulence status of 1440 ascospore-derived strains collected from residues of each cultivar by phenotyping on seedlings. Virulence frequencies were lower in mixtures than in pure stands of the resistant cultivar, as at t2, revealing that the impact of mixtures persisted until the beginning of the next epidemic (t3). The frequency of virulence was lower in the offspring population on the susceptible cultivar and, more surprisingly, the frequency of avirulence on the resistant cultivar increased after sexual reproduction. Our findings highlight two epidemiological processes in mixtures: selection within the local pathogen population between t1 and t2 driven by asexual cross-contamination between cultivars (previous study) and sexual crosses between avirulent and virulent strains between t2 and t3 driven by changes in the probabilities of physical encounters (this study). Mixtures therefore appear to be a promising strategy for the deployment of qualitative resistances, not only to limit the intensity of Septoria tritici blotch epidemics, but also to reduce the erosion of resistances by managing evolution of the pathogen population at a pluriannual scale.

## 1. Introduction

One of the major pillars of crop protection against fungal diseases is the use of resistant cultivars with different sources of resistance. Qualitative resistance genes can confer full protection, by preventing pathogens from infecting, growing on or reproducing in host-plant tissues (Dodds & Rathjen, 2010). However, the deployment of cultivars carrying the same genes at large spatial scales creates strong selective pressure on pathogen populations (McDonald & Linde, 2002). This pressure favours the emergence of ‘virulent’ individuals capable of infecting plants that were hitherto resistant, and increases in the frequency of such individuals (Niks *et al*., 2011). As the frequency of virulent strains increases, the resistance is eroded, and may then be considered to have broken down (Brown & Tellier, 2011). In this context, cereal farming systems are faced with two major concrete problems: (i) the number of resistance genes known and/or available to breeders is limited, and (ii) it may take longer for a gene to be incorporated into new cultivars than for its resistance to be broken down. The current challenge is, therefore, to identify and/or optimise agricultural practices to extend the lifetime of cultivars: in other words, to reinforce the ‘durability’ of their resistance to diseases (Johnson, 1984).

The trade-off between the ‘efficacy’ and ‘durability’ of resistance to plant pathogens has emerged as an important issue in recent years (Mundt, 2018; Rimbaud *et al*., 2018; Cowger & Brown, 2019). The efficacy of strategies for controlling fungal diseases is generally assessed at the scale of the cropping season, because this is the period of interest for a grower, for limiting yield losses due to disease. However, larger time scales should be considered, because a strategy can result in cumulative effects over a period of years, affecting its durability. Ecological studies have highlighted the importance of deciphering the relationship between the season and the ‘off-season’ of the pathogen for epidemiological predictions (Numminen *et al*., 2019). An understanding of this relationship is even more crucial if an important step in the pathogen life cycle occurs predominantly between growing seasons. This is the case for example, for sexual reproduction, which helps to maintain the pathogen population between annual epidemics, while also driving evolutionary processes (Burdon & Laine, 2019). Epidemiological studies have widely highlighted the impact of sexual reproduction on pathogen survival between growing seasons (e.g. Yuen & Andersson, 2013, for *Phytophthora infestans*). Moreover, the intensity of sexual reproduction, the amount of primary inoculum at the beginning of the subsequent epidemic and the size of the pathogen population during this epidemic, for which disease ‘severity’ is a proxy, have been shown to be correlated (e.g. Lo-Pelzer *et al*., 2009, and Fortune *et al*., 2021, for *Leptosphaeria maculans;* Cowger *et al*., 2002; Suffert *et al*., 2018a, for *Zymoseptoria tritici*). The recombination resulting from sexual reproduction also has a qualitative effect on the characteristics of the offspring population relative to the parental population. However, few experimental studies have investigated the impact of sexual reproduction on the frequency of strains displaying virulence against a specific gene within pathogen populations at field scale. Sexual recombination can modify the capacity of a progeny to infect the host on which its parental population originated, and the “beneficial” impact of cropping practices on the pathogen population during the preceding growing season, limiting the increase in virulence frequency, for example.

*Z. tritici* is the ascomycete fungus responsible for *Septoria tritici* blotch (STB), one of the most damaging wheat diseases in temperate countries (Shaw & Royle, 1989; Fones & Gurr, 2015). This pathosystem is a useful model for investigating these issues. *Z. tritici* displays both asexual reproduction, which predominates during the cropping season, and sexual reproduction, which occurs mostly during the interseason period (Suffert *et al*., 2011). During the autumn, the fungus survives on wheat residues, forming sexual fruiting bodies (perithecia) that release wind-dispersed ascospores, which constitute the main source of primary inoculum for the next epidemic season (Sanderson, 1972; Shaw & Royle, 1989). The genetic recombination resulting from sexual reproduction underlies the diversity of the fungal population and the adaptive dynamics of this species (Chen & McDonald, 1996; McDonald & Linde, 2002). As a means of controlling STB, breeders have incorporated *Stb16q*, one of the 21 major resistance genes identified to date (Ghaffary *et al*., 2012; Brown *et al*., 2015), into several cultivars, including cv. Cellule (Florimond Desprez, France). This cultivar was widely grown in France over the last decade (see Fig. 2 in Orellana-Torrejon *et al*., 2021). A few years after the deployment of Cellule (2012), strains capable of overcoming *Stb16q* were detected, and such strains now comprise a substantial proportion of field populations. For example, these strains were estimated to have reached proportions of 0.13 and 0.14 in a trial in the Parisian basin in 2018 and 2019, respectively (Orellana-Torrejon *et al*., 2021).

Growing different cultivars (cultivar mixtures) with complementary sources of resistance in the same field is a promising, well-documented practice for controlling epidemics of plant diseases, such as STB (Cowger & Mundt, 2002; Vidal *et al*., 2017; Kristoffersen *et al*., 2021). In a previous experimental study, we showed that wheat cultivar mixtures can have a significant impact on both the size and composition of *Z. tritici* populations relative to those on pure stands of the cultivar carrying the resistance gene (Cellule), over the course of the growing season (Orellana-Torrejon *et al*., 2021). We established that, in cultivar mixtures, cross-contaminations may occur between cultivars during the cropping season, modifying the dynamics of the pathogen population on a particular cultivar relative to the pure stand. Taking the pure stand of Cellule as a reference, we established that cultivar mixtures could help to slow the dynamics of resistance breakdown.

Our goal in this study was to investigate the consequences of sexual reproduction on the impact of mixtures on the frequency of virulence against a broken-down STB resistance gene within the offspring population constituting the pool of primary inoculum for subsequent epidemics. The challenge was determining whether the evolution of pathogen populations in cultivar mixtures is limited to the scale of an annual epidemic or whether it may continue over a longer period.

## 2. Materials and methods

### 2.1. Overall strategy

In a field experiment divided into microplots, we characterised *Z. tritici* populations in binary mixtures with different proportions of a cultivar carrying the *Stb16q* resistance gene relative to pure stands. The populations studied consisted of strains sampled from diseased leaves at the end of a STB epidemic (t2; see Figure 1 in Orellana-Torrejon *et al*., 2021) and from the corresponding wheat residues after interseason sexual reproduction (t3). We characterised population size based on disease severity at2 in the field and based on the amount of ascospore released from residues at t3 in laboratory conditions. Strains were then phenotyped on both cultivars in greenhouse seedling assays, to compare the aggressiveness and composition of populations based on estimates of the frequency of strains virulent against *Stb16q*.

**Figure 1.**
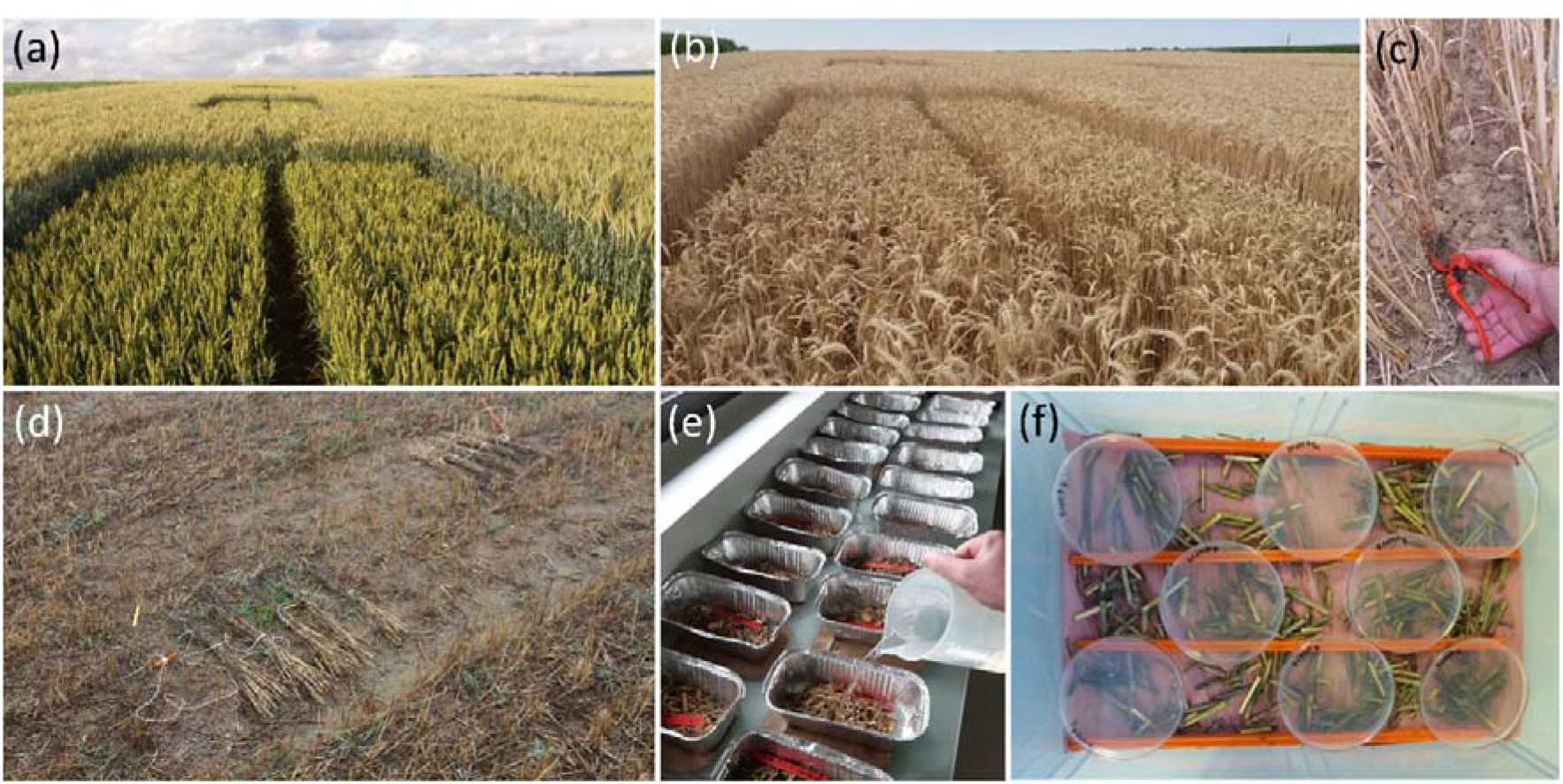
*Zymoseptoria tritici* populations on wheat tissues at the late stage of the epidemic (t2) and after interseason sexual reproduction (t3). (a) Microplot at t2 (11^th^ May 2018). (b) Microplot just before harvest (11^th^ July 2018). (c) Manual collection of the wheat plants. (d) Wheat bundles, separated by cultivar, positioned on the ground of a microplot at t3, i.e. after four months of exposure to weather conditions (26^th^ November 2018). (e) Wetting of wheat residue fragments in laboratory conditions. (f) Petri dishes with PDA medium placed upside down over wheat residue fragments to generate a discharge event and to collect ascospore-derived strains.

**Figure 2.**
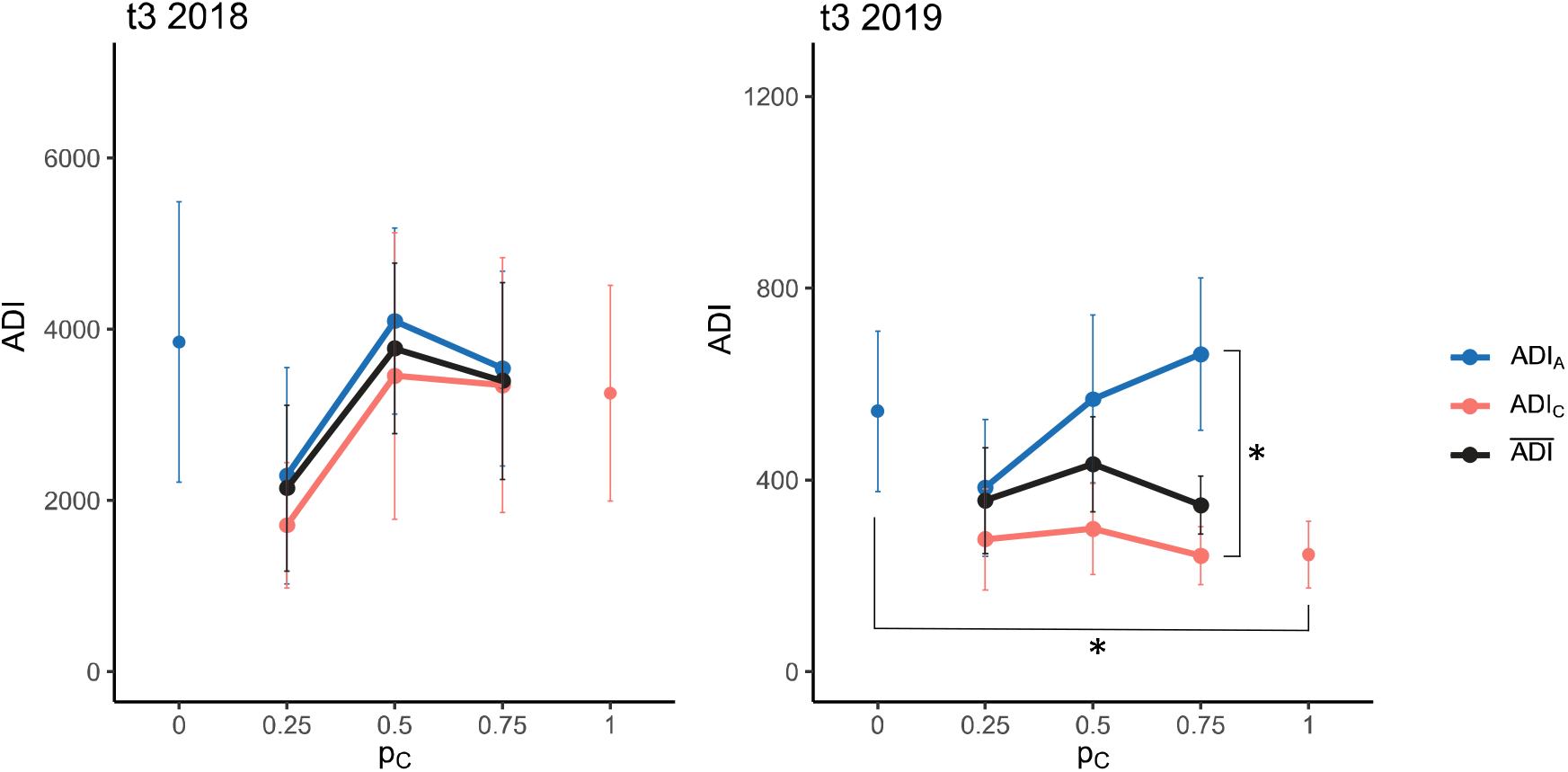
Mean ascospore discharge index, ADI_v_ (mean number of *Zymoseptoria tritici* ascospores collected from 1 g of residues of cultivar v), according to the proportion of cv. Cellule (p_C_) in the canopy at t3, in 2018 and 2019. ADI_v_ was calculated for each cultivar, Apache (ADI_A_, blue solid line) and Cellule (ADI_C_, red solid line), based on six discharge events (two per block). The overall ascospore discharge index 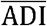 (black solid line) was calculated at the whole-canopy scale as the mean of ADI_A_ and ADI_C_ weighted by the proportions of the cultivars. Bars represent the standard error for the six discharge events. ADI_A_ and ADI_C_ were significantly different (*) only in 2019, in pure stands and C3A1 (ANOVA followed by a Tukey HSD post-hoc test). No other differences between treatments (p_C_) were significant for ADI_A_ or ADI_C_.

### 2.2. Design of the field experiment

This experiment was performed as part of the field trial described in full by Orellana-Torrejon *et al*., 2021. This trial, conducted during the 2017-2018 season and repeated in 2018-2019, was designed to compare the annual dynamics of *Z. tritici* populations in pure stands and in binary mixtures of wheat cultivars, from the early (t1) to the late stage (t2) of an epidemic (Figure 1a). It was divided into three blocks (a, b, and c), each consisting of five microplots treated as follows: a pure stand of wheat cv. Apache (C0A1), a pure stand of wheat cv. Cellule (C1A0), and three binary mixtures, with sowing proportions of Cellule (p_C_) of 0.25, 0.5 and 0.75 (C1A3, C1A1, and C3A1, respectively). Apache (A) was chosen for its moderate susceptibility to STB and Cellule (C) because it carries the *Stb16q* resistance gene for which resistance breakdown has recently been reported.

### 2.3. Management of wheat residues and sexual reproduction

Wheat plants were sampled after their complete natural desiccation, on 10^th^ July 2018 and 16^th^ July 2019 (Figure 1b), over a distance of 2 x 1 m in rows 3 and 4 at the eastern side of each microplot to prevent border effects (Figure S1 in Orellana-Torrejon *et al*., 2021). Before harvest, the plants were cut at a height 5 cm from the ground in each microplot (Figure 1c) and grouped by cultivar, based on the presence (Cellule) or absence (Apache) of awns on the ears. The resulting 24 bundles were stored in a dry place outside the field, for three weeks. Meanwhile, the rest of the plants in the microplots were harvested mechanically on 17^th^ July 2018 and 23^rd^ July 2019 (short cut at a height of 5 cm). On 2^nd^ August 2018 and 14^th^ August 2019, each plant bundle was divided into five smaller bundles, hereafter referred to as sets of ‘residues’, which were deposited on the soil in their microplot of origin, at a location at which most of the other crop debris had been removed by raking. The residues were, thus, exposed to field conditions during the autumn to promote sexual reproduction between *Z. tritici* strains present on the senescent wheat tissues before harvest (Figure 1d). On 26^th^ November 2018 and 10^th^ December 2019, 50 g of decaying plant material (mostly stems and sheaths) was sampled from each set of residues and left to dry in laboratory conditions for one month.

### 2.4. Characterisation of pathogen population size

The intensity of sexual reproduction was estimated by quantifying ascospore production by each set of residues of each cultivar with two simultaneous discharge events per treatment and per block, as described in Suffert *et al*. (2016). For each discharge event, 15 g of residues were cut into 2 cm fragments, soaked in water for 20 min (Figure 1e) and spread on dry filter paper in a box (24 × 36 cm) with the lid left half-open to decrease the relative humidity of the air. Eight Petri dishes (90 mm in diameter) filled with PDA (potato dextrose agar, 39 g · L^-1^) medium were then placed upside down 1 cm above the fragments (Figure 1f). The boxes were placed at 18°C overnight (15 h). The Petri dishes were then closed and incubated at the same temperature in the dark. Six days later, clustered yeast-like colonies resembling cream-coloured convoluted heaps were observed. We counted the number of these colonies by eye, and their identification as *Z. tritici* was checked under a microscope four days after ascospore discharge. It was assumed that each colony resulted from the germination of a single ascospore, and that clusters of colonies appeared just above mature pseudothecia from which several ascospores had been discharged (see Figure 2c in Suffert *et al*., 2016). For each discharge event, we calculated a cumulative ascospore discharge index (ADI; ascospores·g^-1^), defined as the total number of ascospores discharged per gramme of wheat residues. We then calculated the mean of six discharge events, ADI_v_, for each cultivar v (v ∈ [A; C]) and for each treatment. An overall mean ascospore discharge index 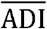 was also calculated at the whole-canopy scale for each treatment, by weighting ADI_v_ by the proportions of the cultivars (p_C_ for Cellule and 1 – p_C_ for Apache), as follows:

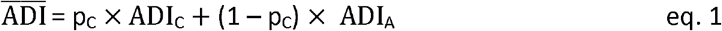

### 2.5. Characterisation of the composition of the pathogen population

#### 2.5.1. Sampling of *Z. tritici* strains from wheat residues

We sampled 30 ascospore-derived strains per cultivar and per microplot (total of 1440 strains, Table 1) from wheat residues. Where possible, the strains were selected from distant clusters of colonies, from as many different Petri dishes as possible, to reduce the probability of sampling the same genotype several times. Each strain was picked with a pin from a single colony and streaked onto a Petri dish containing PDA, which was then incubated at 18°C for five days in the dark. This operation was repeated twice to obtain pure single-genotype strains. Each strain was finally scraped off the dish to obtain a substantial mass of blastopores (150 mg), which was deposited at the bottom of a cryotube and stored dry at −80°C.

**Table 1.** Number of *Zymoseptoria tritici* strains sampled on each cultivar (‘A’ for Apache, ‘C for Cellule) for each treatment (C0A1, C1A3, C1A1, C3A1 and C1A0) after interseason sexual reproduction (t3) following the 2018 and 2019 epidemics, with phenotyping to determine virulence status.

| Year | Stage | C0A1 | C1A3 |  | C1A1 |  | C3A1 |  | C1A0 | Total |
| --- | --- | --- | --- | --- | --- | --- | --- | --- | --- | --- |
|  |  | A | C | A | C | A | C | A | C |  |
| 2018 | t3<sup>1</sup> | 90 | 90 | 90 | 89 | 90 | 90 | 90 | 90 |  |
| 2019 | t3<sup>2</sup> | 88 | 90 | 90 | 90 | 90 | 90 | 90 | 90 |  |

<sup>1</sup> discharged on 26<sup>th</sup> November 2018 from wheat residues collected in the field on 10<sup>th</sup> July 2018
<sup>2</sup> discharged on 10<sup>th</sup> December 2019 from wheat residues collected in the field on 16<sup>th</sup> July 2019

#### 2.5.2. Population phenotyping

Phenotyping data for the *Z. tritici* population sampled at t2 (total of 1440 strains) were obtained and analysed in a previous study (Orellana-Torrejon *et al*., 2021). We characterised the strains sampled at t3 for virulence and aggressiveness on cv. Cellule and Apache seedlings according to the same protocol. This population was phenotyped by splitting it into 14 batches of strains tested in the greenhouse (14 batches of 93 strains and 2 batches of 69 strains). Each batch was balanced to ensure that it contained similar numbers of strains from each cultivar, treatment, and block. Each strain was tested individually on two leaves of three 18-day-old seedlings grown in a pot (six replicates), for each cultivar. For each strain, inoculation was performed with blastospore suspensions (concentration: 10^5^ to 5.0 x 10^5^ spores·mL^-1^) and a cotton bud. All seedlings were then enclosed in transparent polyethylene bags moistened with distilled water for 72 h, to promote infection. The same eight virulent strains as used by Orellana-Torrejon *et al*. (2021) were added to each batch as control strains, to ensure similar conditions for all batches.

Disease severity was assessed by eye 21 days post-inoculation, by determining the percentage of the inoculated leaf surface covered by pycnidia. A strain was considered pathogenic if pycnidia were observed on at least one of the six leaves of Apache. A pathogenic strain was considered virulent (‘vir’) against *Stb16q* if pycnidia were observed on at least one of the six leaves of Cellule, and avirulent otherwise. The frequency of virulent strains *f*_t3, v, vir_ (v ∈ [A; C]) was calculated for each cultivar for each treatment.

We calculated the conditional aggressiveness (AGc; defined as the amount of disease “conditional on the individuals being infected” by McRoberts *et al*., 2003) of each strain sampled on cv. Apache and Cellule at the early (t1) and late (t2) epidemic stages (see Orellana-Torrejon *et al*., 2021) and at t3 (after interseason sexual reproduction; current dataset). For a given strain, AGc was calculated if at least three of the six inoculated leaves on cv. Apache presented pycnidia.

### 2.6. Overall impact of cultivar mixtures on the dynamics of virulent subpopulations after sexual reproduction

We estimated the number of virulent ascospores discharged per gramme of residues (*n*_t3,v,vir_) for each cultivar v (v ∈ [A; C]) as a proxy for the size of the virulent subpopulation at t3. This variable was obtained by multiplying the ascospore discharge index (ADI_v_) by the frequency of virulent strains (*f*_t3,v,vir_) for each cultivar x treatment x block combination. The mean size of the virulent subpopulation at t3 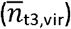 was calculated at the whole-canopy scale for each treatment, as for the overall mean ascospore discharge index (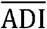, eq. 1).

### 2.7. Statistical analysis

All data analyses were performed with R software (v4.0.2 R Core Team 2012). Shapiro tests (‘shapiro.test’ function) showed that none of the variables studied were normally distributed. We therefore used various approaches to account for the non-normality of the data, depending on the number of replicates available and the actual distribution of the data.

We assessed the effect of treatment on the size of the pathogen population at t3 (i.e. on the intensity of interseason sexual reproduction), by normalising the ascospore discharge index (ADI_v_) data with ordered quantile transformation (*bestNormalize* R package; ‘orderNorm’ function) and performing an analysis of variance (ANOVA; ‘lm’ function). The variance of the overall mean ascospore discharge index 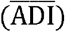 was estimated by calculating error propagation from the standard error of individual variables. We analysed the frequency of virulent strains and the frequency of pathogenic strains during the wheat growing season (t2; t3) with a generalised linear model adapted to a binomial distribution of data (‘glm’ function).

The variance of the three variables described above was partitioned into sources attributable to the following factors and their interactions: cultivar of origin (A; C), treatment (C0A1; C1A3; C1A1; C3A1; C1A0), block (a; b; c), and year (2018; 2019). We also assessed the effect of ‘period’ (t2; t3) on the frequency of virulent and pathogenic strains. We performed a Tukey HSD post-hoc test for multiple pairwise comparisons on estimated marginal means (*emmeans* and *multcomp* R packages; ‘emmeans’ and ‘cld’ functions). The pertinence of the generalised linear models used was checked by calculating the ratio of the residual deviance to the number of degrees of freedom (< 1).

The effect of treatments on the frequency of virulent strains (*f*_t3,v,vir_) and on the frequency of pathogenic strains was assessed with a *χ*^2^ test (‘chisq.test’ function) for contingency tables or with Fisher’s exact test when the expected numbers were small. The effects of treatments on the number of virulent ascospores discharged per gramme of residues for each cultivar (*n*_t3,v,vir_; v ∈ [A; C]) and on the mean size of the virulent subpopulation calculated at the whole-canopy scale 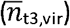 were assessed in a Kruskal-Wallis test (*agricolae* R package, ‘kruskal’ function), selected as the appropriate test for one ANOVA factor with a non-normal distribution. Similarly, the effect of ‘period’ (t1; t2; t3) on conditional aggressiveness means (AGc) was assessed in a Kruskal-Wallis test. Bonferroni correction was used for pairwise comparisons, due to its high stringency.

## 3. Results

### 3.1. Impact of cultivar mixtures on the overall size of pathogen populations on residues after sexual reproduction

The ascospore discharge index (ADI) was 10 times higher in 2018 than in 2019, regardless of cultivar or treatment (p < 0.05, ANOVA; Table S1 and Figure 2). ADI depended on cultivar, without interaction with any other factor. The post-hoc test revealed that ADI_A_ was significantly higher than ADI_C_ in pure stands, and in C3A1, in 2019, highlighting a higher intensity of sexual reproduction in Apache than in Cellule in these two sets of conditions. ADI was highly variable, particularly between blocks. Treatment had a significant effect on the intensity of sexual reproduction for two of the three blocks, but the overall effect, when all three blocks were considered together, was not significant. The effect of treatment on 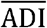 was not significant, at least partly due to the high variability of ADI_A_ and ADI_C_. In 2019, ADI_A_ tended to increase (*p* = 0.038, ANOVA) with the proportion of Cellule in the mixture (p_C_; Figure 2 and Table S1).

### 3.2. Impact of sexual reproduction on the composition of pathogen populations in cultivar mixtures

Three factors had a significant impact on the frequency of virulent strains: year, cultivar and treatment. The change in the frequency of virulent strains between t2 and t3 depended on cultivar (the generalised linear model highlighted a significant period x cultivar interaction; Table S2).

#### 3.2.1. The impact of mixtures persisted after sexual reproduction, on both cultivars

The variation of the frequency of virulent strains (*f*_t3,A,vir_ and *f*_t3,C,vir_) with the proportion of Cellule (p_C_) in the mixture followed the same trend as at the late epidemic stage (t2) for both cultivars (for more details, see Orellana-Torrejon *et al*., 2021). After sexual reproduction (t3), the frequency of virulent strains on Apache was still significantly higher in mixtures than in pure stands, in both years (*χ*^2^ test, *p* < 0.001; Figure 3a and 3b), and it increased with p_C_. At t3 on Cellule, the effect of treatment on the frequency of virulent strains relative to that in the pure stand of Cellule was not significant, except for C1A3 in 2018 (*χ*^2^ test; *p* < 0.001 in 2018 and *p* = 0.208 in 2019; Figure 3c and 3d). The proportion of Cellule (p_C_), thus, had little impact on the frequency of virulent strains on Cellule (*f*_t3,C,vir_), as at t2. Moreover, the difference in the frequency of virulent strains between t2 and t3 was non-significant for almost all treatments and for both cultivars (generalised linear model after post-hoc comparisons). We therefore conclude that the overall effect of mixtures persisted after sexual reproduction, on both cultivars.

**Figure 3.**
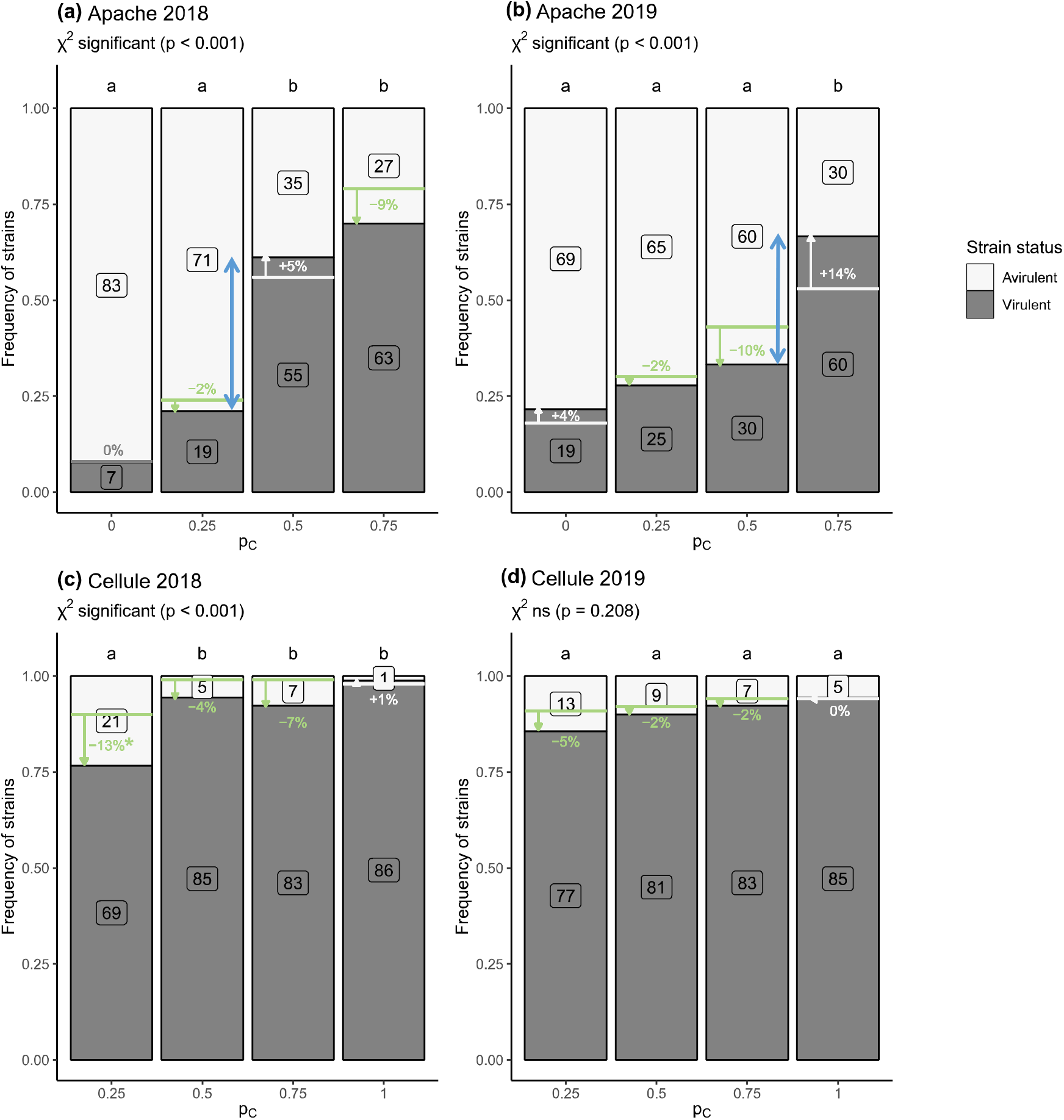
Change in the frequency *of Zymoseptoria tritici* strains considered virulent against *Stb16q* in the populations collected **(a, b)** from cv. Apache (*f*_t3,A,vir_) and **(c, d)** cv. Cellule (*f*_t3,C,vir_) in pure stands (p_C_ = 0) and mixtures (p_C_ = 0.25, 0.5, and 0.75 for C1A3, C1A1, and C3A1, respectively) from t2 (wheat plants at the late epidemic stage) to t3 (wheat residues after interseason sexual reproduction) in 2018 and 2019. The frequencies of virulent and avirulent strains are indicated by the grey and white bars, respectively, and the corresponding numbers appear in boxes. The effect of the treatment (pure stands and cultivar mixtures) on the number of virulent strains was assessed by performing a χ^2^ test with Bonferroni correction for pairwise comparison (letters at the top of each bar). Changes in the number of pathogenic strains from t2 to t3 were calculated for each pc and are represented with simple arrows (white for increases and green for decreases). Blue double arrows highlight the increase in the virulence frequency gap between cultivar proportions from t2 to t3. None of the differences in the number of virulent strains from t2 to t3 within treatments was significant, according to the generalised linear model, except for p_C_ = 0.25 on Cellule in 2018 (*:p< 0.05).

#### 3.3.2. The impact of sexual reproduction (‘period’ effect) on the frequency of virulent strains was dependent on cultivar and treatment (p_C_)

On Apache, the effect of treatment seemed to be emphasized by an inflection in the value of p_C_ at t3, with a substantial increase in the frequency of virulent strains (Figure 3a and 3b). The values obtained were between p_C_ = 0.25 (C1A3) and p_C_ = 0.5 (C1A1) in 2018 (tripling *f*_t3,A,vir_) θnd between p_C_ = 0.5 (C1A1) and p_C_ = 0.75 (C3A1) in 2019 (doubling *f*_t3,A,vir_)o These findings reflect particularly high levels of variation in the frequency of virulent strains after sexual reproduction around this value of p_C_. In particular, fewer virulent strains were found on Cellule in mixtures (regardless of p_C_) at t3 than at t2 in both years, with differences in *f*_t3,C,vir_ ranging from −2% to −13%; however, this difference was significantly only for C1A3 in 2018 (generalised linear model, *p* = 0.020; Figure 3c). This suggests that sexual reproduction led to an overall increase in the frequency of avirulent strains on Cellule in all mixtures (from x1.2 to x7).

### 3.3. Overall impact of mixtures on the dynamics of the virulent subpopulation after sexual reproduction

The virulent subpopulation *n*_t3,v,vir_ was about 10 times larger in 2018 than in 2019, whatever the treatment or cultivar (Figure 4a). After sexual reproduction on Apache, the virulent subpopulation (*n*_t3,A,vir_) was significantly larger in two mixtures (C1A1 and C3A1) than in the corresponding pure stands in 2018 (Kruskal-Wallis, *p* = 0.016; Figure 4a). In 2019, the differences were not significant. The high variability of ADI, the quantitative variable used to calculate *n*_t3,v,vir_, made it impossible to highlight other significant effects. Nevertheless, *n*_t3,A,vir_ increased significantly with the proportion of Cellule (p_C_) in 2018 and there was a similar trend in 2019 (p = 0.016 and p = 0.077, respectively, Figure 4a and 4b). By contrast, on Cellule, differences in *n*_t3,C,vir_ between mixtures and the pure stand were not significant, in either year (Figure 4c and 4d). Finally, the size of the overall virulent subpopulation estimated at whole-canopy scale 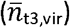 increased with the proportion of Cellule in the mixture, but this trend was not statistically significant (Figure 4e and 4f).

**Figure 4.**
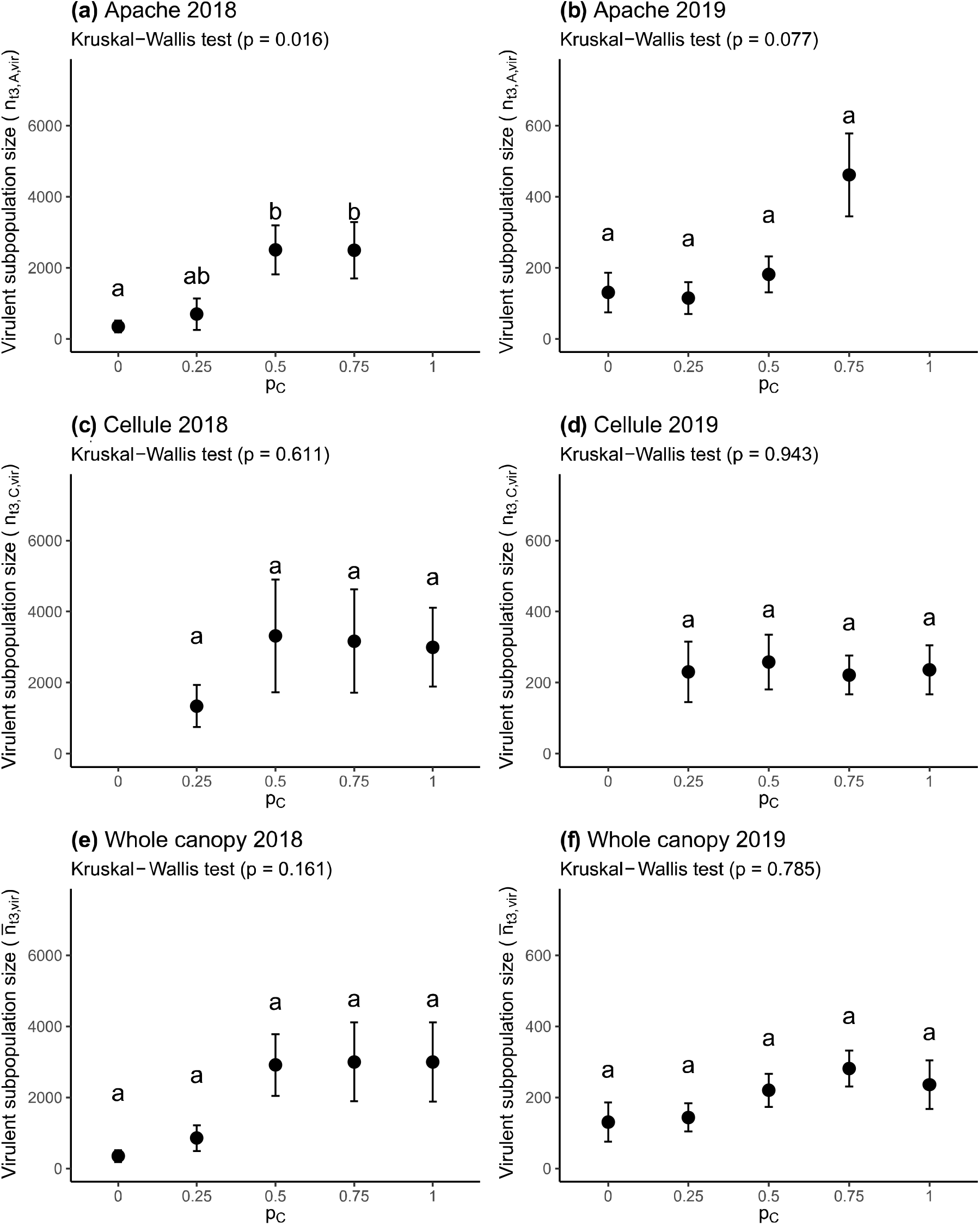
Impact of mixtures on the size of the *Zymoseptoria tritici* virulent subpopulation after sexual reproduction (t3) on wheat residues from each cultivar and at the whole-canopy scale. (a, b) Proxy for the size of the virulent subpopulation sampled from residues of cv. Apache (*n*_t3,A,vir_; number of virulent strains collected from 1 g of residues; black dots) and (c, d) from residues of cv. Cellule (*n*_t3,C,vir_), according to the proportion of Cellule (p_C_) in 2018 and 2019. (e, f) Proxy for the size of the virulent subpopulation calculated at the wholecanopy scale 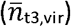. Bars represent the standard error for six replicates for *n*_t3,A,vir_ and *n*_t3,C,vir_, and three replicates for 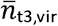. Error propagation was taken into account. The effect of treatment (pure stand or cultivar mixture) on the mean size of the virulent subpopulation was assessed by performing a Kruskal-Wallis test with Bonferroni correction for pairwise comparison.

### 3.4. Pathogenicity and aggressiveness of pathogen populations after sexual reproduction

As at t2, most of the strains sampled from Apache and Cellule residues (> 95%) were pathogenic, causing sporulating lesions on Apache (Figure S1). The proportion of Cellule in the mixture (p_C_) had no significant effect on the frequency of pathogenic strains on either cultivar, except for C3A1 on Apache in 2019 and on Cellule in 2018 (Fisher’s test *p* = 0.016 and *p* = 0.008, respectively, Figure S1). The difference in the frequency of pathogenic strains between t2 and t3 was not significant overall and depended on treatment (p = 0.015 for the interaction between period and treatment; Table S2). The post-hoc comparisons showed that the differences were significant on Apache in C0A1 and C1A3 in 2018 and on Cellule in C3A1 in 2018 and C1A3 and C3A1 in 2019.

The conditional aggressiveness (AGc) of the *Z. tritici* populations sampled at t1, t2, and t3, expressed as the mean percentage sporulating leaf area on cv. Apache in conditions in which at least three of the six inoculated leaves presented pycnidia, was calculated for a total of 3563 strains. AGc decreased significantly after sexual reproduction in 2018, and a similar trend was observed in 2019 (Kruskal-Wallis, p < 0.01 and p = 0.08, respectively; Figure 5). AGc was 18.4% for strains collected at t3, versus 20.4% at t1 and 22.0% at t2 in 2018, and 12.4% for strains collected at t3, versus 16.0% at t1 and 14.4% at t2 in 2019. The differences observed between t1 and t2 were not significant in either year.

**Figure 5.**
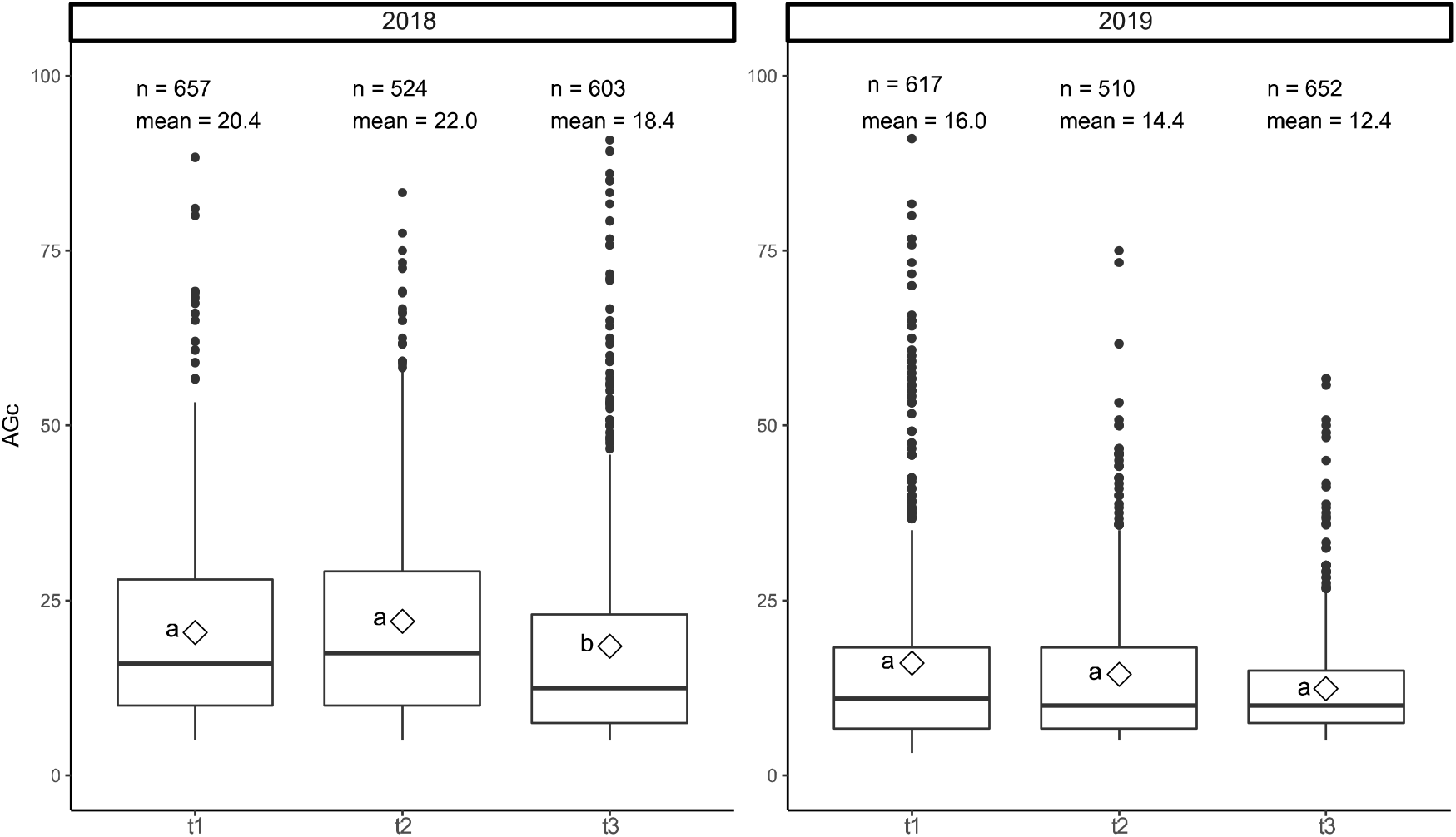
Comparison of the conditional aggressiveness (AGc) of 3563 strains collected on Apache and Cellule in the field trial, by sampling period: early epidemic stage (t1), late epidemic stage (t2) and after interseason sexual reproduction (t3). AGc was calculated as the mean percentage of the leaf area displaying sporulation on cv. Apache in conditions in which at least three of the six inoculated leaves presented pycnidia. For each sampling period, the number of strains (n), the mean (white diamond) and the median (horizontal bar) are indicated. The effect of the sampling period was assessed in a Kruskal-Wallis test (p < 0.01 and *p* = 0.08 in 2018 and 2019, respectively).

## Discussion

This study follows on from a previous study (Orellana-Torrejon *et al*., 2021), in which we highlighted the impact of mixtures of cultivars between the early (t1) and late (t2) stages of the wheat growing season on both the overall size of *Z. tritici* populations (with disease ‘severity’ as a proxy) and their composition (frequency of virulent strains and size of the virulent subpopulation). We show here that sexual reproduction during the interseason period (t3) on wheat residues does not greatly affect the beneficial impact of mixtures over the epidemic season: the selective dynamics displayed during the growing season persisted in the offspring populations. The frequency of virulent strains remained lower in mixtures than in pure stands of Cellule (hereafter referred to as the ‘resistant’ cultivar), indicating that mixtures effectively limited the transmission of virulence against *Stb16q* resistance into the following epidemic season. Mixtures had no detectable impact on the size of offspring populations relative to pure stands, contrary to expectations of a possible decrease in the amount of primary inoculum based on the lower disease severity observed at t2. We discuss here, in greater detail, the novelty and promise of this study, together with its limitations, by distinguishing the effects on each cultivar and proposing explanatory mechanisms.

One of our key findings concerns the impact of sexual reproduction on the composition of the offspring population and the persistence of the overall mixture effect, with differences between cultivars. After interseason sexual reproduction, the increase in the proportion of the susceptible cultivar in the mixture was accompanied by: (i) a decrease in the frequency of virulent strains on the susceptible cultivar (treatment effect; Figure 3a and 3b), consistent with the results obtained late in the growing season (t2) (Orellana-Torrejon *et al*., 2021); and (ii) an increase in the frequency of avirulent strains on the resistant cultivar (treatment effect; Figure 3c and 3d). The dual processes involved in reducing the frequency of virulence in mixtures justified a cultivar-by-cultivar analysis. The effect of the proportion of the cultivar in the mixture was significant for the susceptible cultivar in both years, and for the resistant cultivar only in 2018. This suggests that the intensity of epidemics, which is dependent on climatic conditions in the spring, may account for the observed differences in the magnitude of the mixture effect between years (see the discussion in Orellana-Torrejon *et al*., 2021).

A high proportion of the susceptible cultivar in mixtures decreased the frequency of virulent strains on this cultivar. No significant differences in the frequency of virulent strains on the susceptible cultivar were observed between t2 and t3, for each mixture considered individually. Thus, the selective dynamics established at t2 persisted during the interseason period. The similarity in the frequencies of virulent strains in populations before and after sexual reproduction is consistent with the genetics of *Z. tritici* as a haploid fungus and with Hardy-Weinberg equilibrium, according to which, the allele and genotype frequencies in a population remain constant from generation to generation in the absence of other evolutionary forces. However, for high proportions of the susceptible cultivar, the frequency of avirulent offspring collected on this cultivar increased sharply around a value identified as a potential inflection point (blue double arrows in Figure 3a and 3b). The gap widened from t2 to t3, highlighting an amplification of the effect of cultivar proportion (p_C_), which could be an emerging property of mixtures.

Conversely, mixtures enriched the offspring populations in avirulence through sexual reproduction on the resistant cultivar. The frequency of avirulent strains remained low on the resistant cultivar in mixtures after sexual reproduction (Figure 3c and 3d). This frequency was higher at t3 (11% on average) than at t2 (6% on average), for all mixtures and all years, and this difference was significant for C1A3 in 2018. The frequency of avirulent strains involved in sexual reproduction was actually higher than the frequency estimated at t2 on the basis of sporulating leaf lesion sampling, and was, thus, higher than expected according to Hardy-Weinberg equilibrium. One possible reason for this difference is that several parental avirulent strains may have been present on the resistant cultivar without causing symptoms, thereby escaping sampling at t2. Avirulent strains may be highly abundant on resistant plants due to splash-dispersed pycnidiospores from susceptible plants, especially in mixtures with a high proportion of the susceptible cultivar (low p_C_). Avirulent strains may have participated in sexual reproduction in a cryptic manner, without the generation of asexual fruiting bodies or symptoms. This would be consistent with the larger increase in the frequency of avirulent strains on the resistant cultivar, significantly so for C1A3 in 2018 (10% at t2 vs. 23% at t3), when the epidemic was more intense. This important finding indicates that sexual reproduction led to not only the maintenance of avirulence, but also its enrichment, in the offspring population on resistant plants. The possibility of crossing between an avirulent and a virulent strain on a plant carrying the corresponding resistance gene was demonstrated by Kema *et al*. (2018) for *Stb6*. Two biological phenomena may account for the asymptomatic persistence of one of the two parental strains: (i) the germination of pycnidiospores on the leaf surface, with the ramification and epiphytic growth of hyphae (Fones *et al*., 2017) or proliferation in the form of blastospores (Francisco *et al*., 2019); (ii) colonisation of the substomatal cavities, with blocking in the leaf tissues before growth. It is also possible that avirulent strains successfully penetrated the plant tissues and caused symptoms as ‘stowaways’ co-infecting the plant simultaneously with virulent strains after a ‘systemic induced susceptibility’ (SIS) reaction (Seybold *et al*., 2020). This hypothesis is plausible because avirulent isolates were isolated from 3% (24/719) of the sporulating lesions on the resistant cultivar at t1 (Orellana-Torrejon *et al*., 2021). These hypotheses require experimental testing under semi-controlled conditions.

Another key finding worthy of discussion is the lower aggressiveness of strains collected from wheat residues after sexual reproduction than of those collected from the living plant at the end of the growing season. The mean aggressiveness of the offspring population (t3) was lower than that of both the initial (t1) and late (t2) parental populations (Figure 5), this difference being significant in 2018. This result is of great interest because it is robust, being based on the *in planta* individual phenotyping of a very large number of strains (> 3500), compensating for the small mean size of sporulating areas, as pointed out by Orellana-Torrejon *et al*. (2021). Secondly, it increases our empirical knowledge of the impact of (mal)adaptation to the host on the short-term evolutionary trajectory of local *Z. tritici* populations (Morais *et al*., 2016; Suffert *et al*., 2018b). This difference concerns the parental population (both t1 and t2) sampled on plants, acting as a fitness selective ‘filter’, and also the offspring population consisting of ascospore-producing individuals sampled before they infected new wheat plants (i.e. before they were ‘filtered’). Moreover, more ‘non-pathogenic’ strains were sampled in mixtures after sexual reproduction (t3) than at t2 (Figure S1), confirming that the offspring population was, on average ‘less adapted’ to its host than the parental population. This decrease in aggressiveness after sexual reproduction is consistent with the pluriannual dynamics established in Suffert *et al*. (2018b), even though we did not observe changes in aggressiveness during the growing season, between t1 and t2 (Figure 5). Suffert *et al*. (2018b) found that strains sampled early in the epidemic were fitter than those sampled at the late stage, but no such increase was observed at the start of the next epidemic. The lower aggressiveness of the offspring population highlighted here confirms the existence of a trade-off between the seasonal (epidemic) and interseason (interepidemic) periods.

The impact of mixtures on the size of offspring populations is an issue meriting further investigation. We did not highlight an impact on the overall size of the *Z. tritici* offspring populations (t3) relative to pure stands. The size of the offspring population was determined by counting the number of ascospores discharged per 1 g of wheat residues (ADI_v_), as a proxy for the intensity of sexual reproduction. The variability of the results, due to the limitations of this method (Suffert & Sache, 2011; Suffert *et al*., 2016, 2018b), together with a significant block effect (Table S1), probably limited our ability to detect an effect in this trial. However, we were able to establish that the intensity of sexual reproduction was significantly higher in 2018 than in 2019 (Figure 2). The intensity of epidemics may account for the observed year effect, as weather conditions were more favourable in spring 2018 than in 2019 for disease development through the splash dispersal of pycnidiospores, resulting in a higher disease severity in both the mixtures and the pure stands of the resistant cultivar. We also found that sexual reproduction was significantly more intense on the susceptible cultivar than on the resistant cultivar, in pure stands and in C3A1 in 2019. This finding is consistent with previous demonstrations of a positive correlation between the intensity of sexual reproduction in several ascomycete pathogens and disease severity during the growing season, as shown by Cowger *et al*., 2002, and Suffert *et al*., 2018a. There are two possible reasons for the absence of such a correlation in this study (Figure S2). First, disease severity was quite low (< 30%) at t2, whereas Suffert *et al*. (2018a) reported a positive correlation only in semi-controlled conditions in which STB severity ranged from 40% to 80%. Second, the residues from which the intensity of sexual reproduction was estimated in this study consisted of all leaf layers and stems, whereas disease severity at t2 was estimated from the last three leaf layers only. This difference in the plant material used may have had a major impact, as the wheat tissues playing the greatest role in sexual reproduction are located in the central part of the plant (ascospore production maximal between 25 and 35 cm above the ground; see Figure 4 in Suffert *et al*., 2018a).

We assessed the effect of sexual reproduction on the frequency of virulent strains in each mixture, but we were unable to evaluate the effect on the change in overall population size and in the size of the virulent subpopulation from t2 to t3 directly. Indeed, the sizes of the two populations were assessed by means of two different proxies: disease severity, expressed as the percentage of the leaf are displaying sporulation (assumed to be correlated with the number of pycnidiospores), and the number of ascospores ejected. We highlighted differences in the size of virulent offspring subpopulations (Figure 4): (i) these populations were 10 times larger in 2018 than in 2019; (ii) they were also larger on susceptible cultivar in mixtures than in pure stands. Given the variability observed with the method for estimating the intensity of sexual reproduction, it seems more relevant to focus on “the frequency of virulence” than on “population size”. Our conclusions on the effects of mixtures should be considered with caution, because there may be an overall trade-off between size and composition in populations (Orellana-Torrejon *et al*., 2021).

In conclusion, the results for the epidemic (Orellana-Torrejon *et al*., 2021) and interseason (the current study) periods reveal that two epidemiological processes can account for the efficacy of cultivar mixtures for limiting the virulent fraction of a local *Z. tritici* population over the course of the complete pathogen life cycle in field. The first of these processes is the short-term selection that occurs within a local pathogen population, driven by cross-contaminations between cultivars with splash-dispersed pycnidiospores (the composition of which, in virulent strains, depends on the cultivar of origin). The second process results from changes in the probabilities of physical encounters between avirulent and virulent strains at the late epidemic stage, as consequence of the first process. We found that this second process is probably based on two mechanisms for decreasing the frequency of virulence in mixtures relative to the resistant pure stands, depending on the cultivar considered in the mixture: (i) a decrease in the frequency of virulent strains on the susceptible cultivar; (ii) more unexpectedly, an enrichment in avirulent strains on the resistant cultivar. These empirical findings confirm that cultivar mixtures are a promising strategy for resistance deployment, not only to limit the intensity of STB epidemics (Kristoffersen *et al*., 2020), but also to manage the erosion of *Stb* genes through effects on virulence frequencies in local *Z. tritici* populations. This study provides experimental elements for further investigation, by modelling approaches, of the potential of cultivar mixtures as a durable resistance deployment strategy relative to other strategies, as assessed by Rimbaud *et al*.(2018), for example. In estimations of the impact of key factors, it will be crucial to consider the nature and number of resistance genes, the proportion of each cultivar in the mixture, and the frequency of virulences (and their combination, if any) in the local population (Vidal *et al*., 2020; Kristoffersen *et al*., 2021). An understanding of the interactions between these factors will clearly require modelling approaches, but also further experimental fungal biology studies in controlled conditions to characterise the abovementioned mechanisms in greater detail. The potential impact of resistance deployment strategies, including cultivar mixtures — which may be beneficial, but also detrimental over the years — remain to be fully assessed. Our conclusions were drawn at field scale, but it should clearly be borne in mind that the impact of mixtures is diluted at greater spatial scales, when considering inoculum exchanges between fields planted with different cultivars (e.g. Fabre *et al*., 2015).

## Acknowledgements

We thank Christophe Montagnier (INRAE Experimental Unit, Thiverval-Grignon, FR), Sandrine Gélisse, Nathalie Retout, Laurent Gerard, Nicolas Lecutier, Auriane Pinton, Miléna Bonnard, Jérôme Lageyre, Martin Willigsecker and Benjamin Boudier (INRAE BIOGER, Thiverval-Grignon, FR) for technical assistance. We thank Dr. Agathe Ballu and Dr. Anne-Lise Boixel for their help and advice with the statistical analysis. We thank Dr. Julie Sappa for her help correcting our English.

## Data availability statement

The data that support the findings of this study are freely available from the INRAE Dataverse online data repository (https://data.inrae.fr/) at xxxxxxx.

---

**Table S1.** Impact of the various experimental factors (year, cultivar, treatment, and block) on the intensity of *Zymoseptoria tritici* sexual reproduction estimated by the ascospore discharge index (ADI; mean number of ascospores collected per 1 g of residues), according to ANOVA. The main effects and their interactions were tested and summarised on the basis of their p values (F test).

| Factor | Sum of square | Degrees of freedom | F statistic | F test | Significance |
| --- | --- | --- | --- | --- | --- |
| year | 41.683 | 1 | 155.168 | < 0.001 | *** |
| cultivar | 2.099 | 1 | 7.815 | 0.007 | ** |
| treatment | 2.454 | 3 | 3.045 | 0.038 | * |
| block | 13.541 | 2 | 25.204 | < 0.001 | *** |
| year x cultivar | 0.483 | 1 | 1.798 | 0.186 | ns |
| year x treatment | 0.175 | 3 | 0.217 | 0.884 | ns |
| cultivar x treatment | 0.327 | 3 | 0.406 | 0.749 | ns |
| year x block | 7.174 | 2 | 13.354 | < 0.001 | *** |
| cultivar x block | 0.037 | 2 | 0.069 | 0.934 | ns |
| treatment x block | 5.698 | 6 | 3.536 | 0.006 | ** |
| year x cultivar x treatment | 1.023 | 3 | 1.270 | 0.295 | ns |
| year x cultivar x block | 0.040 | 2 | 0.074 | 0.929 | ns |
| year x treatment x block | 3.084 | 6 | 1.914 | 0.098 | . |
| cultivar x treatment x block | 2.831 | 6 | 1.757 | 0.128 | ns |
| year x cultivar x treatment x block | 1.187 | 6 | 0.737 | 0.623 | ns |

**Table S2.** Impact of the various experimental factors (year, period, cultivar, treatment, and block) on the number of strains virulent against *Stb16q* (i.e., causing sporulating lesions on cv. Cellule) and pathogenic strains (i.e., causing sporulating lesions on cv. Apache) in *Zymoseptoria tritici* populations collected on Apache and Cellule in the field trial, estimated with a generalised linear model. The main effects and their interactions were tested and summarised on the basis of their p values (F test). The models were fitted to retain only the significant interactions.

| Factor | LR $\chi^2$ | Degrees of freedom | $\chi^2$ probability | Significance |
| --- | --- | --- | --- | --- |
| Number of virulent strains |  |  |  |  |
| year | 4.53 | 1 | 0.033 | * |
| period | 2.47 | 1 | 0.116 | ns |
| cultivar | 582.37 | 1 | < 0.001 | *** |
| treatment | 296.34 | 4 | < 0.001 | *** |
| block | 1.85 | 2 | 0.396 | ns |
| period x cultivar | 7.02 | 1 | 0.008 | ** |
| year x treatment | 43.18 | 4 | < 0.001 | *** |
| cultivar x treatment | 5.38 | 2 | 0.068 | . |
| period x block | 6.01 | 2 | 0.050 | * |
| period x cultivar x block | 9.36 | 4 | 0.053 | . |
| Number of pathogenic strains |  |  |  |  |
| year | 17.31 | 1 | < 0.001 | *** |
| period | 0.20 | 1 | 0.652 | ns |
| cultivar | 0.66 | 1 | 0.418 | ns |
| treatment | 7.02 | 4 | 0.135 | ns |
| block | 9.29 | 2 | 0.010 | ** |
| period x treatment | 12.40 | 4 | 0.015 | * |
| treatment x block | 25.88 | 8 | 0.001 | ** |
| year x period x treatment | 16.80 | 8 | 0.032 | * |
| year x cultivar x treatment | 16.72 | 5 | 0.005 | ** |
| period x cultivar x treatment | 10.53 | 3 | 0.015 | * |
| year x treatment x block | 21.44 | 10 | 0.018 | * |

**Figure S1.**
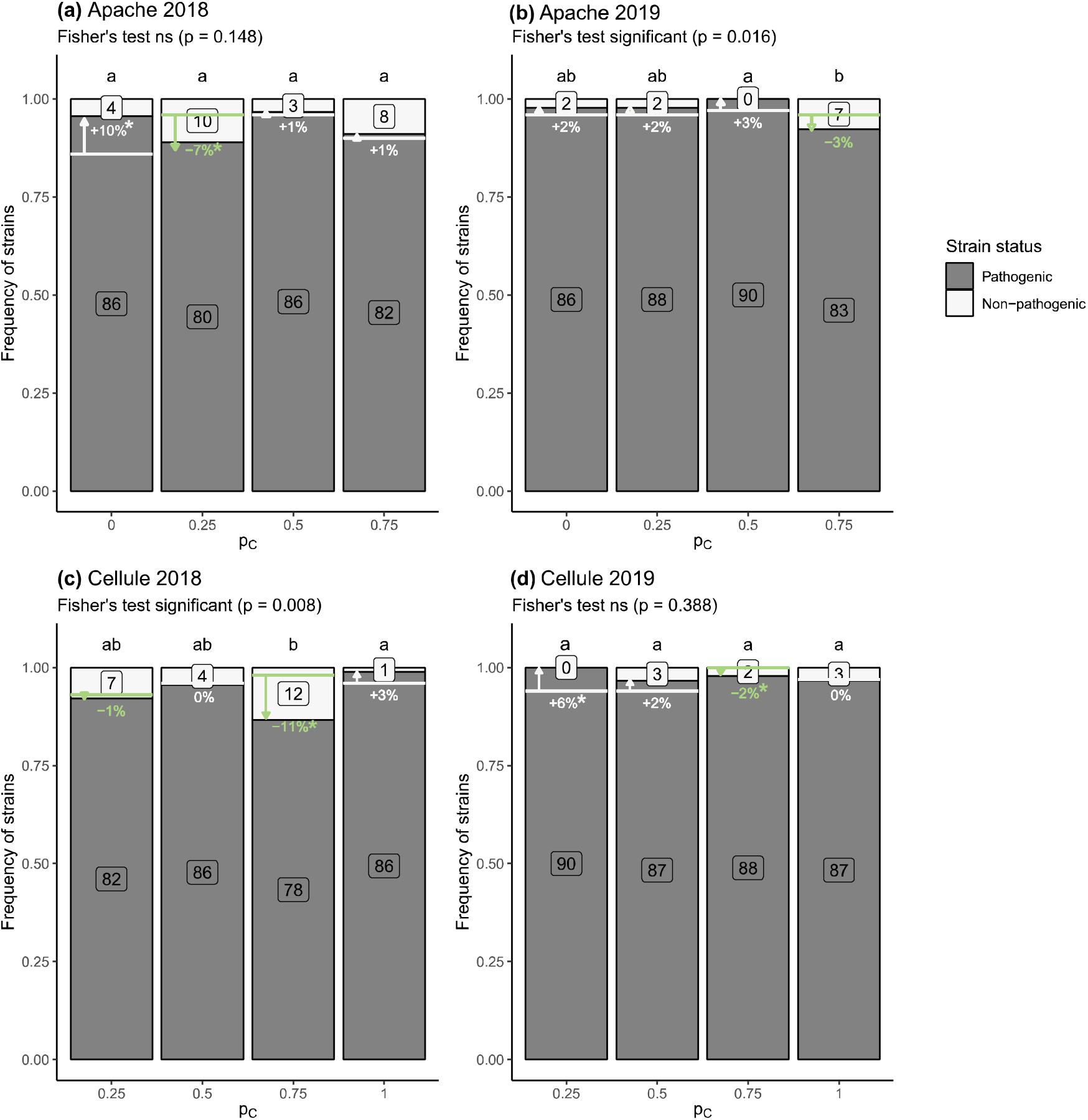
Change in the frequency *of Zymoseptoria tritici* strains considered pathogenic in the populations collected **(a, b)** on cv. Apache and **(c, d)** cv. Cellule in pure stands (p_C_ = 0) and mixtures (p_C_ = 0.25, 0.5, and 0.75 for C1A3, C1A1, and C3A1, respectively) from t2 (wheat plants at the late epidemic stage) to t3 (wheat residues after interseasonal sexual reproduction) in 2018 and 2019. The frequencies of pathogenic and non-pathogenic strains are indicated with grey and white bars, respectively, and the corresponding numbers appear in boxes. The effect of treatment (pure stands and cultivar mixtures) on the number of pathogenic strains was assessed by performing a χ^2^, or Fisher’s exact test when the expected numbers were small, with Bonferroni correction for pairwise comparisons (letters at the top of each bar). Changes in the number of pathogenic strains from t2 to t3 were calculated for each p_C_ and are represented with arrows (white for increases and green for decreases). Significant differences in the number of pathogenic strains from t2 to t3 within each treatment are indicated according to the results for the generalised linear model (*: *p* < 0.05).

**Figure S2.**
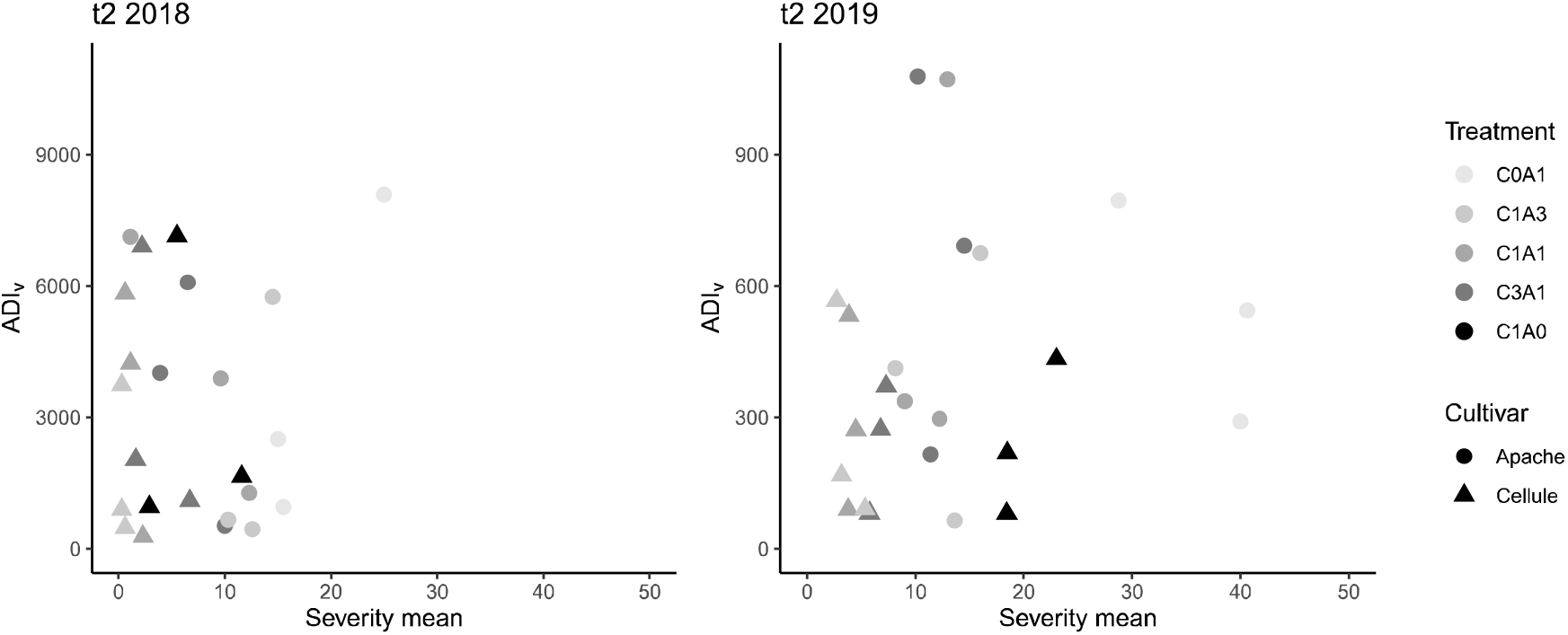
Relationship between the intensity of *Zymoseptoria tritici* sexual reproduction (ADI_v_) on wheat residues during the interseason period (t3) and disease severity at the late stage of the epidemic (t2). Disease severity was calculated in the field as the mean percentage of the leaf area covered with pycnidia, for flag leaves in 2018 and for the second and third leaves from the top in 2019. The intensity of sexual reproduction was estimated by calculating the ascospore discharge index (ADI; mean number of ascospores collected per 1 g of residues), for cv. Apache (red symbols) and for cv. Cellule (blue symbols), for each block. Symbols correspond to the different treatments (C0A1 and C1A0 for pure stands of Apache and Cellule, respectively; C1A3, C1A1 and C3A1 for the mixtures).

